# Using machine learning models to predict oxygen saturation following ventilator support adjustment in critically ill children: a single center pilot study

**DOI:** 10.1101/334896

**Authors:** Sam Ghazal, Michael Sauthier, David Brossier, Wassim Bouachir, Philippe Jouvet, Rita Noumeir

## Abstract

Clinicians’ experts in mechanical ventilation are not continuously at each patient’s bedside in an intensive care unit to adjust mechanical ventilation settings and to analyze the impact of ventilator settings adjustments on gas exchange. The development of clinical decision support systems analyzing patients’ data in real time offers an opportunity to fill this gap. The objective of this study was to determine whether a machine learning predictive model could be trained on a set of clinical data and used to predict hemoglobin oxygen saturation 5 min after a ventilator setting change. Data of mechanically ventilated children admitted between May 2015 and April 2017 were included and extracted from a high-resolution research database. More than 7.10^5^ rows of data were obtained from 610 patients, discretized into 3 class labels. Due to data imbalance, four different data balancing process were applied and two machine learning models (artificial neural network and Bootstrap aggregation of complex decision trees) were trained and tested on these four different balanced datasets. The best model predicted SpO_2_ with accuracies of 76%, 62% and 96% for the SpO_2_ class “< 84%”, “85 to 91%” and “> 92%”, respectively. This pilot study using machine learning predictive model resulted in an algorithm with good accuracy. To obtain a robust algorithm, more data are needed, suggesting the need of multicenter pediatric intensive care high resolution databases.

## Introduction

In case of respiratory failure, mechanical ventilation supports the oxygen (O_2_) diffusion into the lungs and the carbon dioxide (CO_2_) body removal. As an expert in mechanical ventilation cannot reasonably be expected to be continuously present at the patient’s bedside, specific medical devices aimed to help in ventilator settings adjustments may help to improve the quality of care. Such devices are developed using either algorithms based on respiratory physiology/medical knowledge that adapt ventilator settings in real time based on patients’ characteristics but are not accurate enough to be used widely in clinical practice, especially in children [1, 2]; or physiologic models that simulate cardiorespiratory responses to mechanical ventilation settings modifications but none was validated for this indication [3]. The above-mentioned models all share the limitation of not being suited to learn from ever-growing sets of clinical research data, and potentially improve their performances. To overcome this drawback, another avenue is the development of algorithms using artificial Intelligence to provide caregivers with support in their decision-making tasks. In this study, we assessed machine learning methods to predict transcutaneous hemoglobin saturation oxygen (SpO_2_) of mechanically ventilated children after a ventilator setting change using a high resolution research database.

## Materials and Methods

This study was conducted at Sainte-Justine Hospital and included the data collected prospectively between May 2015 and April 2017 of all the children, age under 18 years old, admitted to the Pediatric Intensive Care Unit (PICU) who were mechanically ventilated with an endotracheal tube. Patients’ data were excluded if the patient was hemodynamically unstable defined as 2 or more vasoactive drugs delivered at the same time (ie., epinephrine, norepinephrine, dopamine or vasopressin) or with an uncorrected cyanotic heart disease defined by no SpO_2_ > 97% during all PICU stay. All the respiratory data from included patients were extracted from the PICU research database [4], after study approval by the ethical review board of Sainte-Justine hospital (number 2017 1480).

## Data extraction

To determine the data that will be extracted for each child, an item generation was conducted by three physicians (PJ, MS, DB). The resulting items are presented in Fig 1 within their sources, means of extraction and a schematic of the main components of the study. The predictive SpO_2_ value was the SpO_2_ 5 minutes after a change of a ventilator setting. The delay of 5 min corresponded to the shortest period of time to reach a steady state after modification of a ventilator setting [5].

**Fig 1.**
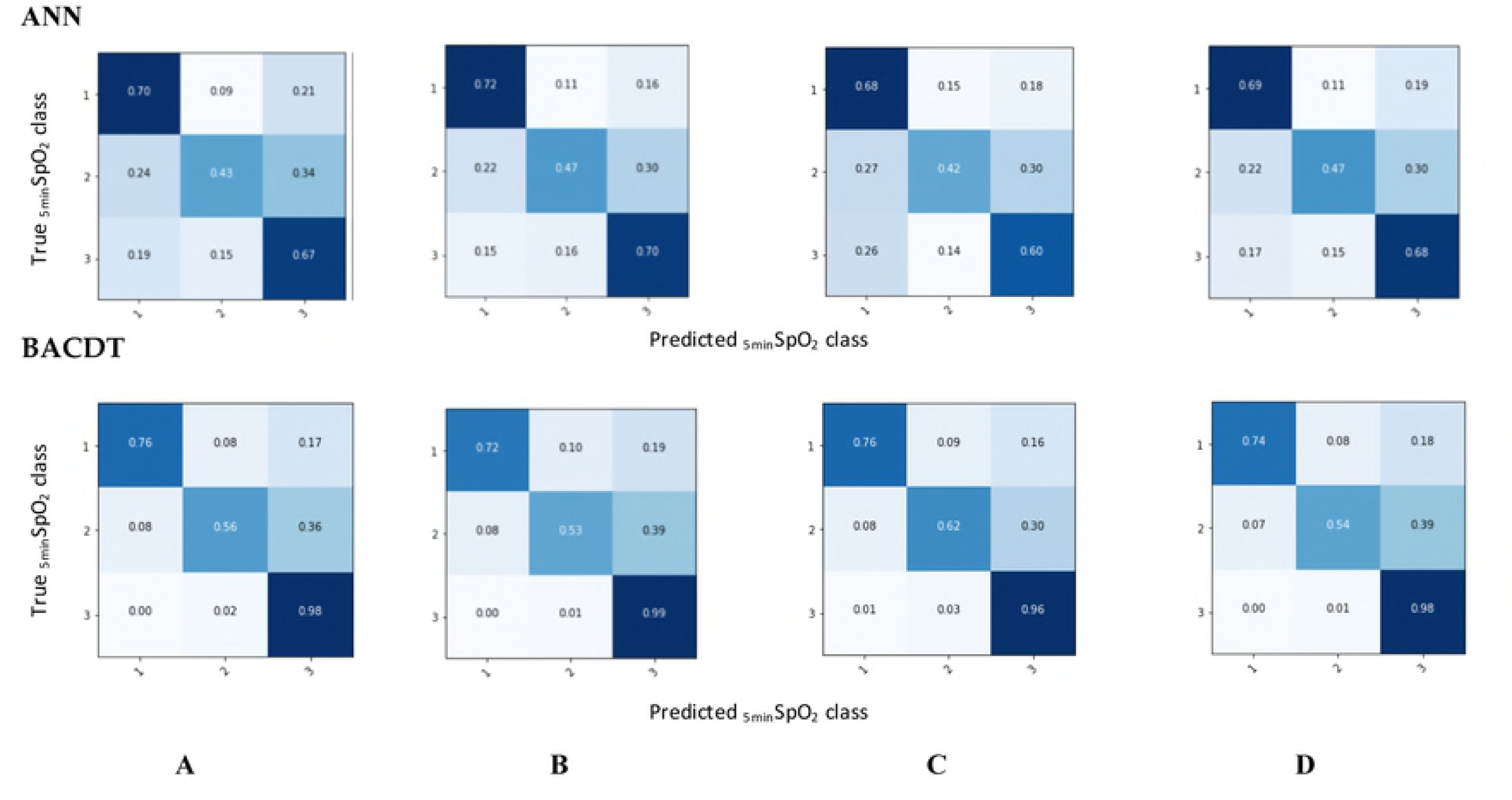
Schematic description of the analysis process and items involved. EMR: electronic Medical Record, FiO_2_: inspired fraction of Oxygen, Vt: tidal volume, PEEP: Positive end expiratory pressure, PS above PEEP: pressure support level Above PEEP, PC above PEEP: pressure control level above PEEP, MVe: expiratory minute volume, I:E Ratio: inspiratory time over expiratory time, Measured RR: respiratory rate measured by the ventilator, PIP: positive inspiratory pressure ie maximal pressure measured during inspiration. _5min_SpO_2_: SpO_2_ observed 5 min after PEEP, FiO_2_, tidal volume, PS above PEEP, PC above PEEP change, ML: machine learning, ANN: artificial neural network, BACDT: Bootstrap aggregation complex decision trees.

## Data formatting

The data extracted from the research database needed: (1) to remove erroneous data due to disconnection of the patient from the ventilator or the monitor, or due to transient interventions such as suctioning; (2) to remove the rows at which no ventilator setting variables was modified; (3) to adapt data format for classifier training. The methodology to format the data is described in S1 file.

## Data categorization

SpO_2_ levels at 5min were classified into three categories (Table 1). The thresholds were selected according to clinical value: a SpO_2_ < 92% is a target to increase oxygenation in mechanically ventilated children [6]. The critical level of 85% SpO_2_ is used as an alarm of severe hypoxemia in intensive care [7].

**Table 1:** Definition of SpO_2_ class labels specifications

| SpO<sub>2</sub> classification | SpO<sub>2</sub> range (%) | Rows number (n) |
| --- | --- | --- |
| 1 | < 84 | 17,112 |
| 2 | 85 to 91 | 29,869 |
| 3 | 92 to 100 | 729,746 |

## Data balancing

The data analysis showed a severe imbalance with most SpO_2_ at 5min above 92%. This is logical as caregivers want to maintain SpO_2_ in normal range during child PICU stay. In such condition, the classifier learns the majority class label (class 3) (Table 1) but doesn’t learn the minority class labels (class 1 and 2) [8]. The data balancing process aims to allow the classifier to learn from all class equally. The data balancing process used in this study included a combination of down-sampling and up-sampling techniques: to balance the three classes of the data involved, a down-sampling of the SpO_2_ class 3 using TOMEK algorithm [9] and an over-sampling of SpO_2_ class 1 and 2 using Synthetic Minority Oversampling Technique (SMOTE) [10] were performed.

The creation of synthetic data points by SMOTE can be formulated as follows:

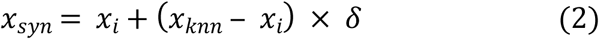

In equation (2), *x*_*syn*_ represents the synthetic data point. The variables *x*_*i*_ and *x*_*knn*_ are respectively the original instance, and the nearest neighbor data point which is randomly picked among the *k* nearest neighbors. The random number *δ* is generated in [0,1] to determine the position of the created synthetic data point along a straight line joining the original data point *x*_*i*_ and its chosen nearest neighbor *x*_*knn*_.

To study which data balancing method provided the more accurate algorithm, four datasets were produced via four different balancing procedures, involving different combinations of data balancing techniques (Fig 2).

**Fig 2.**
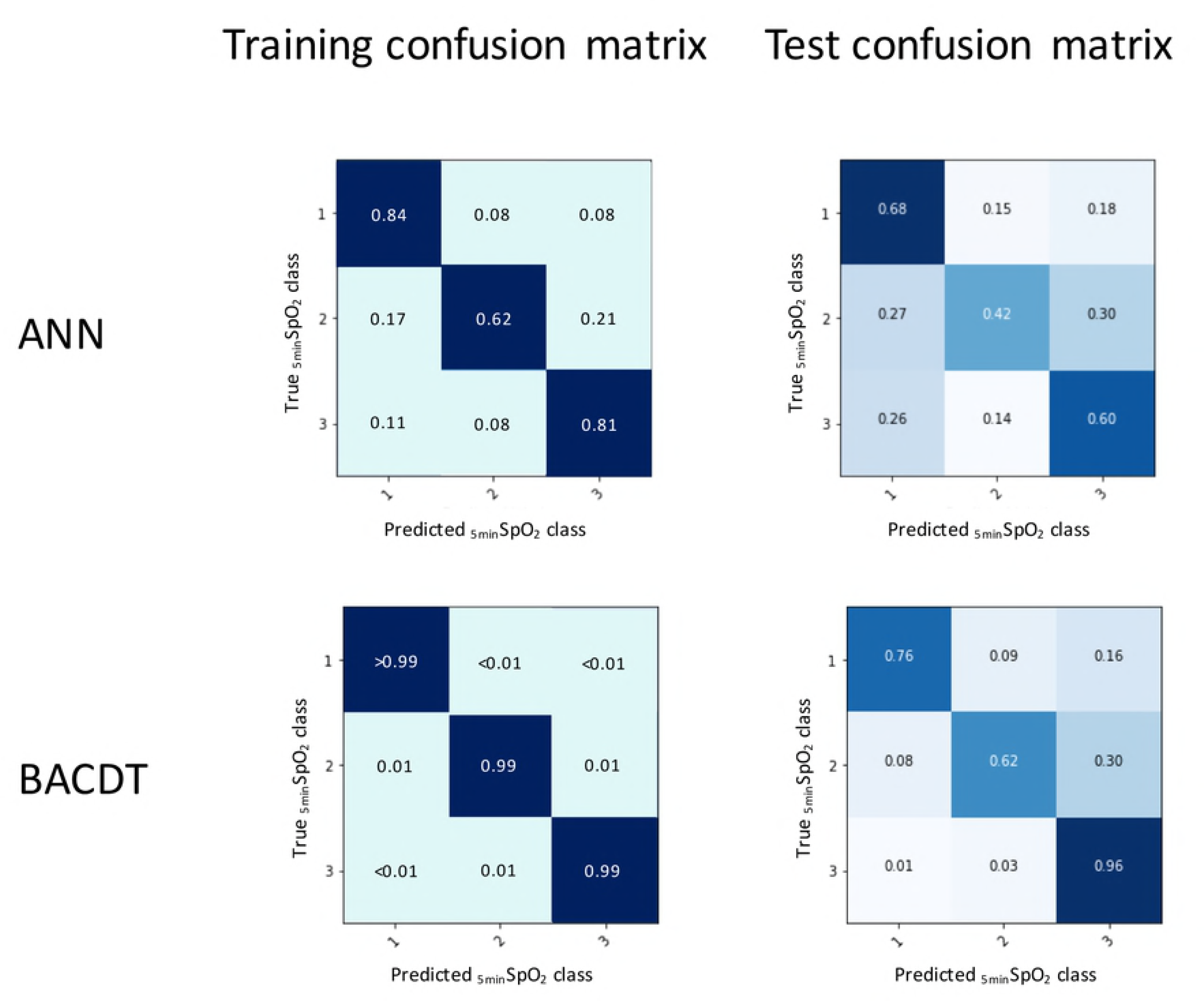
Descriptions of the four balancing procedures.

### Predicted SpO_2_ Classification

To identify the best machine learning classification method, we tested two classification models: artificial neural network and bagged complex decision trees, on the four balanced datasets.

### Artificial Neural Network (ANN)

Once the data has been pre-processed, a machine learning predictive model was trained on a sub-set of labeled training data. The model is then used to predict the target variable values on a testing subset where the class labels are hidden. We used Artificial Neural Networks (ANN) to make predictions of the SpO_2_ variable, based on the values of other variables of interest. Through the function approximation that the ANN performs, it is possible to make predictions of SpO_2_ variable, based on the input data.

The ANN is learned from training data, using the backpropagation algorithm [11] and is tested on a test set made of the remaining rows of data to validate the generalization of the model. The learning algorithm runs through all the rows of data in the training data set and compares the predicted outputs with the target outputs found in the training data set. The weights are adjusted via supervised learning, in a manner to minimize the error of predicted SpO_2_ vs target SpO_2_. The process is repeated until the error is minimized.

The ANN classifier was implemented through cycles of forward propagation followed by backward propagation through the network’s layers. The backpropagation algorithm is used for performance optimization.

For a given number of classes K > 2, the cross-entropy error can be formulated as shown in eq. 3, where (***W*_*i*_**)_*i*_ is the matrix of weights between the neuron layers, *r*_*i*_ is the target value. *y*_*i*_ is the value generated by the ANN, ie., its output.

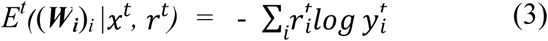

The outputs of the ANN are:

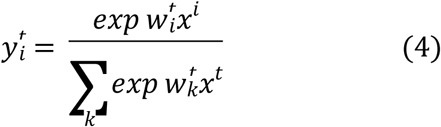

Using stochastic gradient-descent (SGD) for error minimization, the update rule for the ANN weights is:

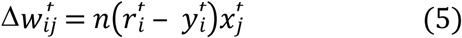

In equation 5, *η* is the learning rate, which, when SGD is used, decreases as the error is minimized. During ANN training, each observation, comprised of an input vector and a target output, is denoted (***x***^*t*^, ***r***^*t*^), with ***r***^*t*^ ϵ (“1”, “2”, “3”). The reason why the cross-entropy (eq. 3) is used instead of the Least Square Error (LSE) is to avoid long periods of training, due to the ANN going through stages of slow error reduction.

### Bootstrap aggregation of complex decision trees

Bootstrap aggregating (acronym: bagging) was proposed by L Breiman in 1994 to improve classification by combining classifications of randomly generated training sets [12]. Bagging allows for the creation of an aggregated predictor via the use of multiple training sub-sets taken from the same training set. Let **(*Ti*)** denote the replicate training sub-sets bootstrapped from the training set ***T***. These replicate sub-sets each contain *N* observations, drawn at random and with replacement from ***T***. For each of these sub-sets of *N* observations, a prediction model, or classifier, is created. The computational model we used for bagging was complex decision trees. This means that, for each bootstrapped sub-set of training data, a complex decision tree is trained and thus a classifier is created. If *i = 1, …, n*, then *n* classifiers are created through the bagging process.

A decision tree is a flowchart computational model which can be used for both regression, as well as classification problems. Paths from the root of the tree to its various leaf nodes go through decision nodes in which decision rules are applied in a recursive manner, based on values of input variables. Each path represents an observation (***X***, *y*) = (*x*_*1*_*, x*_*2*_*, x*_*3*_*, …, x*_*n*_*, y*), where the label assigned to the target *y* is given in the leaf node, at the end of the path, ie., classification [13].

In the aim of maximizing the model’s generalization capability during the training process, the Bagged Complex Trees’ performance is tested via *k*-fold cross-validation. A value *k* = 10, which is common practice, was used in this study. The training using *k*-fold cross-validation is carried out as described in Fig 3.

**Fig 3.**
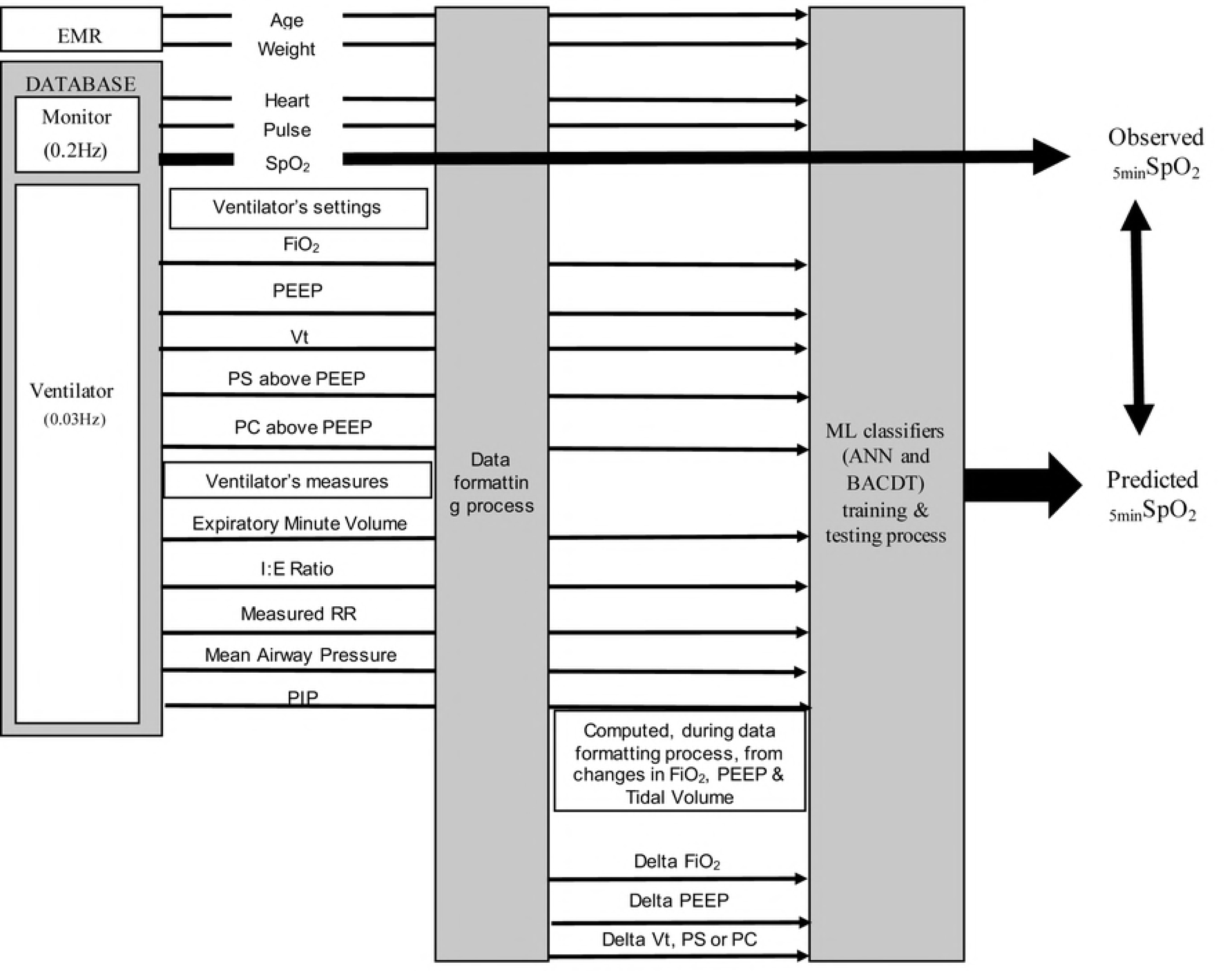
*k*-fold cross-validation.

The mathworks Matlab R2016b Machine Learning toolbox was used for the creation of the ensemble of Bagged complex trees model.

### Assessment of the performances of the classifiers

We evaluated the performances of the classifiers based on the metrics including testing confusion matrix, average accuracy, precision, recall and F score [14] with a _5min_SpO_2_ prediction expected above 0.9 for each class.

### Precision

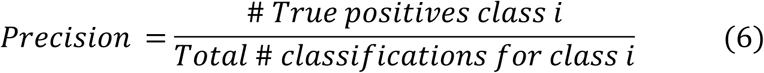

The *Precision* (eq. 6) is the ratio of all correct classifications for class *i* to all instances labeled as class label *i* by the model. In a non-normalized confusion matrix, this would mean dividing the number of instances classified in class label *i* by the total of instances in column *i.*

### Recall

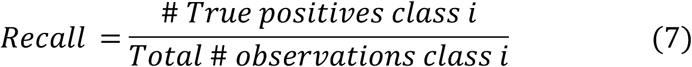

Recall is the ratio of the number of instances classified in class label *i* to the number of true class *i* labels. In a non-normalized matrix, this would require dividing the number of instances classified in class label *i* by the total of row *i*

### F-score

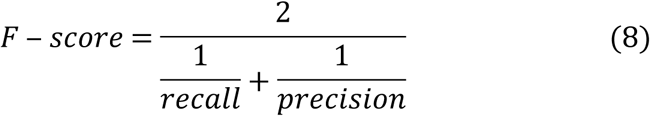

The F-score provides a single measure of classification performance of the model used.

## Results and discussion

We developed and assessed the performances of two machine learning classifiers on four different balanced datasets to predict SpO_2_ at 5 min after a ventilator setting change (*ie* FiO_2_, PEEP, Vt/Pressure), in 610 mechanically ventilated children. In Fig 4 and Table 2, we report the performances of these two classifiers. Using the classification performance metrics, the bagged trees classifier trained on dataset #3 (see Fig 2) has yielded the best classification performance on the test sets (Table 2). The confusion matrix of the whole bagged trees shows that SpO_2_ at 5 min could correctly predict in **76%** of class “1” data, **62%** of class “2”, and **96%** of class “3” (Fig 4). This huge variation in classification performances of the three class labels can be explained by the large variation in the numbers of observations available for each of the class labels in the initial dataset that has limited the machine learning (Table 1).

**Table 2.** Performance of artificial neural networks (ANN) and bootstrap aggregation of complex decision trees (BACDT) classifiers for SpO_2_ prediction at 5 min following a ventilator setting change. Avg/total: average accuracy of total classification values. In italics is the performance of the best predictive model obtained among the eight tested.

| Balanced datasets | 5minSpO<sub>2</sub> class | ANN |  |  | BACDT |  |  |
| --- | --- | --- | --- | --- | --- | --- | --- |
|  |  | Precision | Recall | F-score | Precision | Recall | F-score |
| Dataset 1 | 1 | 0.12 | 0.70 | 0.21 | 0.80 | 0.76 | 0.78 |
|  | 2 | 0.16 | 0.43 | 0.23 | 0.61 | 0.56 | 0.59 |
|  | 3 | 0.96 | 0.67 | 0.79 | 0.97 | 0.98 | 0.97 |
|  | Avg/total | 0.88 | 0.65 | 0.73 | 0.94 | 0.94 | 0.94 |
| Dataset 2 | 1 | 0.09 | 0.72 | 0.16 | 0.77 | 0.72 | 0.74 |
|  | 2 | 0.09 | 0.47 | 0.16 | 0.57 | 0.53 | 0.55 |
|  | 3 | 0.98 | 0.70 | 0.81 | 0.98 | 0.99 | 0.98 |
|  | Avg/total | 0.93 | 0.69 | 0.78 | 0.96 | 0.97 | 0.97 |
| Dataset 3 | 1 | 0.16 | 0.68 | 0.25 | *0.80* | *0.76* | *0.78* |
|  | 2 | 0.26 | 0.42 | 0.33 | *0.67* | *0.62* | *0.65* |
|  | 3 | 0.92 | 0.60 | 0.72 | *0.95* | *0.96* | *0.96* |
|  | Avg/total | 0.80 | 0.58 | 0.65 | *0.91* | *0.91* | *0.91* |
| Dataset 4 | 1 | 0.09 | 0.69 | 0.16 | 0.80 | 0.74 | 0.77 |
|  | 2 | 0.12 | 0.47 | 0.19 | 0.58 | 0.54 | 0.56 |
|  | 3 | 0.97 | 0.68 | 0.80 | 0.98 | 0.98 | 0.98 |
|  | Avg/total | 0.92 | 0.67 | 0.76 | 0.96 | 0.96 | 0.96 |

**Fig 4.**
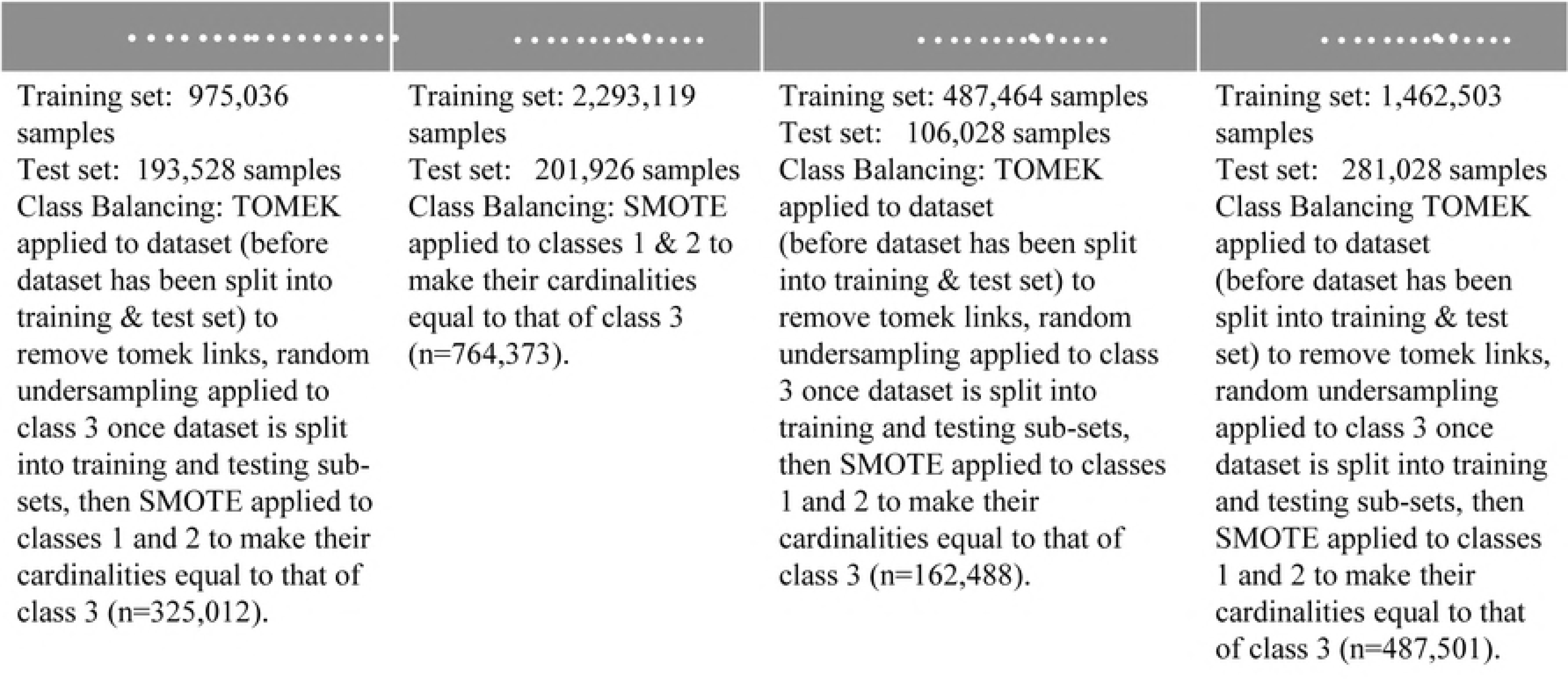
Artificial neural network (ANN) and bootstrap aggregation of complex decision trees (BACDT) test confusion matrices. The darker colors represent higher levels of accuracy. A: balanced dataset 1, B: balanced dataset 2, C: balanced dataset 3, D: balanced dataset 4 (see Fig 2).

For the artificial neural network, the variation of the number of hidden layers and number of neurons per hidden layer did not seem to have a significant effect on the model’s classification performance (Table 3). As for the Bagged complex trees, the variation of the number of complex trees did not yield significant changes in classification performance (Table 4).

**Table 3.**
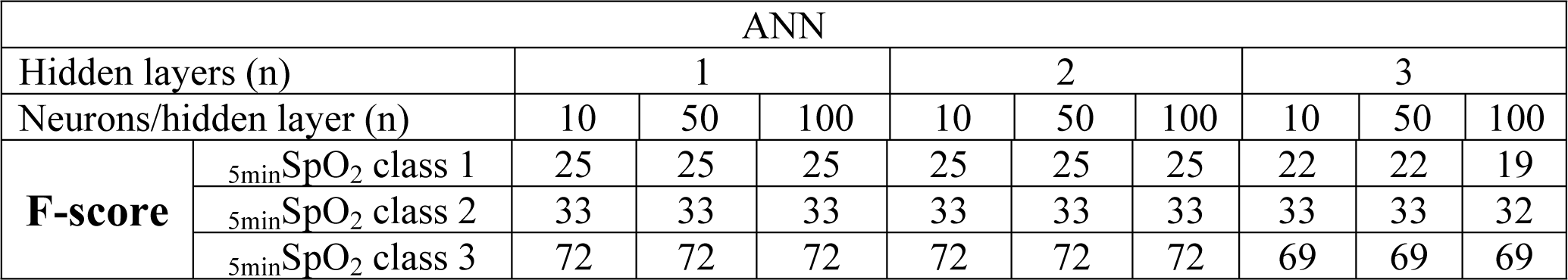
Absence of impact on performance of the increase of neurons and hidden layers for artificial neural network (ANN). Example of the performance assessed by the F score on the balanced dataset 3 (see fig 2)

| ANN |  |  |  |  |  |  |  |  |  |  |
| --- | --- | --- | --- | --- | --- | --- | --- | --- | --- | --- |
| Hidden layers (n) |  | 1 |  |  | 2 |  |  | 3 |  |  |
| Neurons/hidden layer (n) |  | 10 | 50 | 100 | 10 | 50 | 100 | 10 | 50 | 100 |
| F-score | $_{5min}SpO_2$ class 1 | 25 | 25 | 25 | 25 | 25 | 25 | 22 | 22 | 19 |
| | $_{5min}SpO_2$ class 2 | 33 | 33 | 33 | 33 | 33 | 33 | 33 | 33 | 32 |
| | $_{5min}SpO_2$ class 3 | 72 | 72 | 72 | 72 | 72 | 72 | 69 | 69 | 69 |

**Table 4.** Absence of impact on performance of the number of complex trees for bootstrap aggregation of complex decision trees (BACDT). Example of the performance assessed by the F score on the balanced dataset 3 (see Fig 2)

|  |  | BACDT |  |
| --- | --- | --- | --- |
|  |  | n = 30 | n=50 |
| **F-score** | $_{5min}SpO_2$ class 1 | 78 | 78 |
| | $_{5min}SpO_2$ class 2 | 65 | 65 |
| | $_{5min}SpO_2$ class 3 | 96 | 96 |

In agreement with previous studies regarding bagging being a better method for medical data classification, tree Bagging fared better than the artificial neural network used in this study [12]. It is noteworthy however that the gaps in performance results between the training and testing confusion matrices are relatively higher in the case of bagged trees model than in that of the artificial neural network (Fig 5). This seems to indicate that, although the bagged trees model was capable of learning very well from the data, there’s still room for improvement in the generalization. The SMOTE algorithm is designed in such a way that should theoretically not affect the generalization of the trained model. In cases of extreme data imbalance, however, as is the case in this study, the over-sampling within the data space of a given minority class label, used for increasing the cardinality of the class label’s set, is also likely to be extreme. This may render the data space of this class relatively dense with respect to the rest of the data, made up of real data points of the studied patient sub-population. This may potentially explain the classification model’s relatively poor generalization for _5min_SpO_2_ class “1” and “2” with respect to the generalization for _5min_SpO_2_ class “3”. Also, since SMOTE generates synthetic data points by interpolating between existing minority class instances, it can obviously increase the risk of over-fitting when classifying minority class labels, since it may duplicate minority class instances. The fact that the training confusion matrix shows extremely high classification performances for the minority _5min_SpO_2_ class “1” and “2”, as opposed to those shown in the testing confusion matrix, suggests that the over-sampling of the minority _5min_SpO_2_ class using SMOTE could have caused some overfitting for these classes, but this would have to be further investigated.

**Fig 5.**
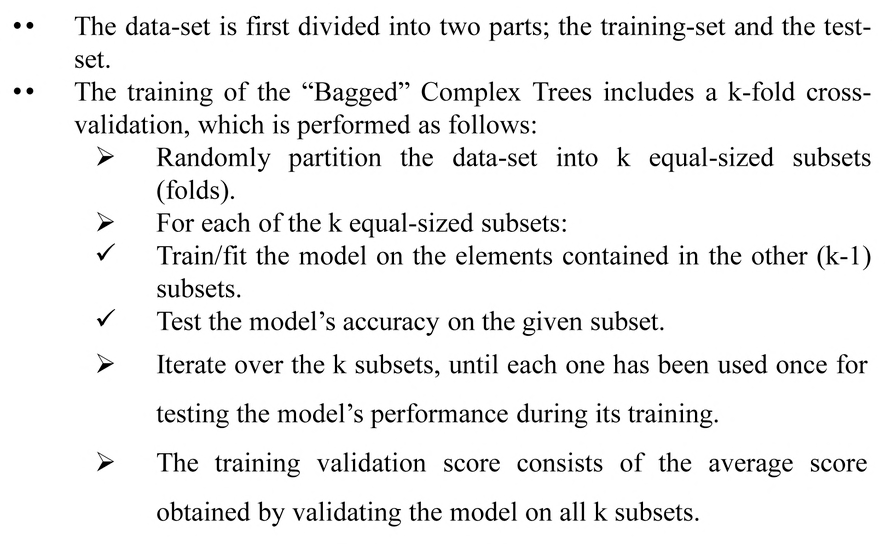
Training and testing confusion matrices of artificial neural networks (ANN) and bootstrap aggregation of complex decision trees (BACDT) classifiers for SpO_2_ prediction at 5 min following a ventilator setting change.

The strengths of this study include a large clinical database of mechanically ventilated children used with more than 7.10^5^ rows. In a recent similar study in PICU, 200 patients were included with 1.15.10^3^ rows [15]. However, the volume of data is clearly insufficient. To use such machine learning predictive models, the pediatric intensive care community needs to combine multicenter high resolution database. In addition, children data could be pooled to neonatal and adult intensive care data, when possible, such as MIMIC III database [16]. The other strength is the process used to transform the data into a usable format and to correct a variety of artifacts present (S1 file). In health care, there is a significant interest in using clinical databases including dynamic and patient-specific information into clinical decision support algorithms. The ubiquitous monitoring of critical care units’ patients has generated a wealth of data which presents many opportunities in this domain. However, when developing algorithms domains, such as transport or finance, data are specifically collected for research purposes. This is not the case in healthcare where the primary objective of data collection systems is to document clinical activity, resulting in several issues to address in data collection, data validation and complex data analysis [17]. As detailed in S1 file, a significant amount of effort is needed, when data have been successfully archived and retrieved, to transform the data into a usable format for research.

This study has several limitations. The limited row number reduced the SpO_2_ classification for machine learning predictive model to three clinically relevant classes. SpO_2_ is a continuous variable and the use of three class is probably insufficient, especially when high SpO_2_ range is suggested as potentially harmful [18, 19]. Instead of the classification model, the next step could be to test regression models’ performance. SpO_2_ was predicted at 5min after ventilator setting change, a clinically relevant delay. However, the delay between ventilator setting change and oxygenation steady state is not well defined and vary from 1 to 71 minutes according to the parameter set (FiO_2_, PEEP or other parameters that change mean airway pressure) and clinical conditions studied [15, 20, 21]. This needs further research and probably more sophisticated clinical decision support systems using machine learning predictive models should consider these factors. Finally, we excluded hemodynamic unstable patients using a treatment criteria (≥ 2 vasoactive drugs infused) because this condition decreases pulse oximeter reliability [22, 23]. The validation and electronic availability of reliable markers of hemodynamic instability in children such as plethysmographic variability indices could be helpful [24].

## Conclusion

This pilot study using machine learning predictive model resulted in an algorithm with good accuracy. To obtain a robust algorithm with such a method, more data rows are needed, suggesting the need of multicenter pediatric intensive care high resolution databases.

## Acknowledgments

We would like to thank Mr. Redha Eltaani for his support in all tasks related to data access at Ste-Justine Hospital. This work was supported by the Natural Sciences and Engineering Research Council of Canada (NSERC), by the Institut de Valorisation des Données (IVADO), by grants from the “Fonds de Recherche du Québec – Santé (FRQS)”, the Quebec Ministry of Health and Sainte Justine Hospital.

## Supporting information

**S1 File: Data formatting process**

